# SLFN11 restricts escape from telomere crisis to prevent alternative lengthening of telomeres

**DOI:** 10.1101/2025.09.10.675313

**Authors:** Sandra Segura-Bayona, Valerie Borel, Tyler H Stanage, Marija Maric, Melanie Walter, Rafaela A Oliveira, Shudong Li, Aurora I Idilli, Martina Peritore, Graeme Hewitt, Maarten Hekkelman, Daniel M Snell, Sam T Jones, Chris Cheshire, Areda Elezi, Probir Chakravarty, Richard Mitter, Anna Mikolajczak, Harriet E Gee, Emma Nye, Roderick J O’Sullivan, Aatur D Singhi, Thijn R Brummelkamp, Anthony J Cesare, Simon J Boulton

## Abstract

The tRNA nuclease SLFN11 is epigenetically silenced in ∼50% of treatment-naive tumours and is the strongest predictor of chemoresistance but why it is frequently inactivated in cancer is unknown. To acquire immortality, cancer cells can activate alternative lengthening of telomeres (ALT), typically accompanied by *ATRX* loss. Here, we implicate SLFN11 in sensing telomere replication stress, triggering eradication of *ATRX* deficient cells prior to ALT establishment. Whereas progressive telomere shortening of cells lacking telomerase and *ATRX* leads to telomere crisis and cell death, *SLFN11* loss confers tolerance to PML-BLM dependent ALT intermediates, permitting emergence of ALT survivors. We propose that during tumorigenesis *SLFN11* inactivation is selected as means to tolerate endogenous replication stress following telomere crisis, leading to the development of therapy resistant tumours before treatment.

## Main

The viral restriction factor SLFN11 (*1*) is a putative tumour suppressor that is lost in ∼50% of all cancers due to promoter hypermethylation. The status of *SLFN11* expression in cancer is the best predictive biomarker of chemotherapy response in patients (*2–4*). While its silencing has been suggested to be a result of selective pressure exerted by DNA damage treatment (*5*), why it is silenced at such a high frequency in treatment-naive cancers is unknown. Also, the *SLFN11* status in cancer cell lines does not correlate with cancer types caused by oncogenic viruses (*3*). The challenge that *SLFN11* silencing poses to cancer treatment underscores the urgent need to better understand mechanisms of tumour evolution that lead to naive chemoresistance.

Tumorigenesis requires cancer cells to acquire a telomere maintenance mechanism to counteract telomere shortening. Progressive telomere erosion eventually triggers two independent anti-tumour mechanisms: replicative senescence and telomere crisis (*6, 7*). To solve the end replication problem and attain replicative immortality, cancers either re-express the reverse transcriptase telomerase to extend telomere repeats (*8*) or induce ALT, which relies on telomere recombination (*9*). ALT-positive cancers account for ∼15% of all malignancies and exhibit specific features not typically observed in telomerase-expressing or primary cells. ALT+ telomeres are often heterogeneous in length while presenting with high rates of telomere sister chromatid exchanges (*10*). Mechanisms of break-induced replication operate in ALT to mediate telomere extension (*11*), requiring recombination factors such as the BLM-TOP3A-RMI1-2 (BTR) complex (*12–14*), which colocalise with clustered telomeres within a subset of subnuclear promyelocytic leukaemia (PML) bodies, termed ALT-associated PML bodies (APBs) (*12, 14–17*). The presence of ALT-specific recombination by-products such as extrachromosomal partially single-stranded telomeric (CCCTAA)_n_ DNA circles, referred to as C-circles (*18*), and native single-stranded (ss) telomeric DNA foci (ssTeloC) (*14*) represent established markers of ALT activity. Loss-of-function mutations in the chromatin remodeller α-thalassemia/mental retardation syndrome X-linked protein (*ATRX*) is the most common identifiable alteration associated with ALT (*19, 20*). Paradoxically, *ATRX* inactivation alone is insufficient for ALT establishment (*20–25*) suggesting the requirement of additional unidentified events that synergise with *ATRX* loss to acquire and sustain ALT.

Here, we report *SLFN11* loss as a key driver event that allows *ATRX* and telomerase deficient cells to escape telomere crisis and become ALT. Our findings establish SLFN11 as a molecular gatekeeper that restricts ALT activation through direct sensing of telomere replication stress and triggering apoptosis. Loss of this safeguarding mechanism permits the emergence of immortalized, therapy-resistant cell populations.

## SLFN11, PML and BTR activities are detrimental to ATRX-null cells

Since *ATRX* loss alone is insufficient for ALT establishment (*20–25*), we set out to identify mechanisms that cooperate with *ATRX* loss to allow ALT. To this end, we established isogenic *ATRX* knockout (KO) in telomerase-positive chronic myeloid leukaemia-derived diploid eHAP and fibrosarcoma HT1080 cells (fig. S1, A and B). When compared to the ALT+ U2OS cell line, *ATRX*-null cells exhibited low levels of ALT hallmarks, including APBs and native ssTeloC foci; a condition we define here as an ALT ‘primed’ state (Fig. 1A and fig. S1, C to F). Acquisition of a broader spectrum of ALT features including generation of extrachromosomal C-circles and enhanced ssTeloC signal was triggered in *ATRX*-null cells upon exposure to chemical inducers of replication stress, such as hydroxyurea (HU) (Fig. 1A and fig. S1, F to I). However, *ATRX*-null cells rapidly lost viability upon exogenous replication stress (Fig. 1B and fig. S1J). Complementation with wild-type *ATRX* attenuated C-circle production (fig. S1, H and K), indicating that *ATRX* loss does not switch irreversibly to an ALT-primed state.

**Fig. 1.**
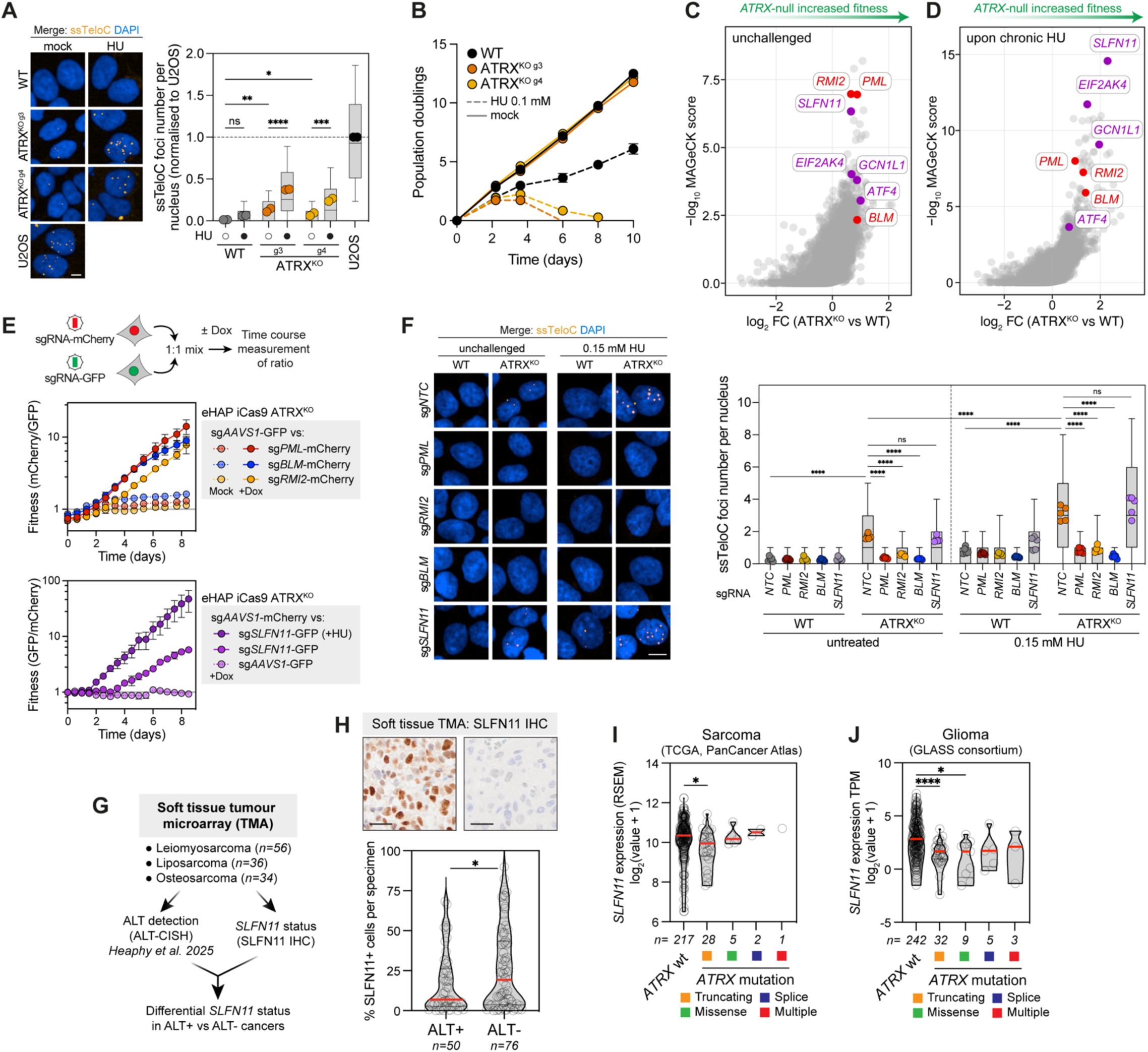
SLFN11, PML and BTR activities are detrimental to *ATRX*-null cells. **A**, Left: Representative native FISH ssTeloC micrographs in eHAP cells ± 24 h 0.15 mM HU. Scale bar: 10 µm. Right: Normalised ssTeloC foci number per nucleus. Box plots represent individual nuclei with interquartile range (IQR), horizontal line indicates median, and whiskers extend to 10^th^ and 90^th^ percentile. Data points outside this range are not shown. Coloured dots represent mean (n=2 biological replicates with minimum 800 nuclei per replicate, one-way ANOVA). **B**, Growth curves in eHAP iCas9 cells ± 0.1 mM HU. Data are mean ± SD (n=3 biological replicates). **C-D**, Volcano plot showing log transformed MAGeCK sgRNA score and fold change for enriched sgRNAs in *ATRX*-null vs WT cells, unchallenged (C) or upon 0.06 mM 10-day HU treatment (D). For more details, see Supplementary Table 3. **E**, Two-colour competitive growth assays in *ATRX*-null cells transduced with the indicated sgRNAs ± doxycycline (Dox). For *AAVS1* vs *SLFN11*, ± 0.06 mM HU in the presence of Dox. Data are mean ± SEM (n=3 biological replicates). **F**, Left: Representative native FISH ssTeloC micrographs in eHAP cells ± 24 h 0.15 mM HU. Scale bar: 10 µm. Right: Number of ssTeloC foci per nucleus. Box plots represent individual nuclei with IQR, horizontal line indicates median, and whiskers extend to 10^th^ and 90^th^ percentile. Data points outside this range are not shown. Coloured dots represent mean (n=5 biological replicates with 500 nuclei as minimum average per replicate, one-way ANOVA). **G**, Schematic of methodology for determination of SLFN11 protein expression in soft tissue cancers. ALT status was determined in (*29*) by chromogenic in situ hybridization (CISH). **H**, Human soft tissue tumour FFPE sections stained with SLFN11. Percentage of cells SLFN11-positive per specimen were quantified by QuPath (unpaired t test with Welch’s correction). **I-J**, *SLFN11* mRNA expression data from human sarcoma or glioma cohorts, classified by *ATRX* mutation. Violin plots with median in red (one-way ANOVA with Šídák’s multiple comparisons correction).

To identify driver events that cooperate with *ATRX* loss to allow ALT establishment, we leveraged genome-scale CRISPR/Cas9 screens in the parental (WT) and *ATRX*-null diploid eHAP cells. Using the MAGeCK algorithm, we analysed enrichment of sgRNA counts relative to parental cells in unchallenged conditions (Fig. 1C) or upon chronic 0.06 mM HU treatment (Fig. 1D). *SLFN11*-targeting sgRNAs were the most significantly enriched in *ATRX*-null cells, conferring a marked increase in cellular fitness upon chronic HU treatment and scoring top-6 hit in unchallenged conditions (Fig. 1, C and D). Counterintuitively, sgRNAs targeting the ALT drivers *PML* and the BTR complex were also enriched in our screen (Fig. 1, C and D), suggestive of cytotoxicity caused by ALT-like by-products. To validate the synthetic viable interactions, we undertook competitive growth assays in *ATRX*-null cells with sgRNAs targeting *PML*, *BLM*, *RMI2, SLFN11* or the safe harbour locus *AAVS1*. In unchallenged conditions, sgRNAs targeting *PML*, *BLM, RMI2* or *SLFN11* caused a growth advantage over those targeting *AAVS1* (Fig. 1E), implying that SLFN11, PML and BTR activities are detrimental to cell viability in ALT-primed *ATRX*-null cells.

To determine how drivers of *ATRX*-null cells fitness impact ALT hallmarks, we examined ssTeloC and C-circles in inducible knockouts of *SLFN11*, *PML* or the BTR complex. Unchallenged *ATRX*-null cells presented with ALT hallmarks that were exacerbated when subjected to HU treatment (Fig. 1A and fig. S1, F to I) and these were completely suppressed by loss of PML or BTR activity (Fig. 1F and fig. S2, A to D). These data are in line with evidence implicating the BLM helicase in the generation of ALT-specific extrachromosomal ssDNA species through excessive telomere lagging strand displacements (*17, 26*). Notably, ssTeloC signal and C-circles remained unaffected in *ATRX*-null cells irrespective of *SLFN11* status (Fig. 1F and fig. S2, C and D), indicating the loss of *SLFN11* enables *ATRX*-null cells to tolerate hallmarks of ALT. Given SLFN11’s role in blocking replication forks upon DNA damage (*27, 28*), we next asked whether SLFN11 is responsible for the sensitivity to exogenous replication stress in *ATRX* deficient cells. Depletion of *SLFN11* completely alleviated sensitivity of *ATRX*-null cells to heightened replicative stress caused by HU or aphidicolin treatment despite not alleviating the actual DNA damage signalling (Fig. 1E and fig. S2, E to G), suggesting a role for SLFN11 in replication stress tolerance. Collectively, these data reveal that ALT activity in *ATRX*-null cells is toxic. Pro-survival mechanisms include either suppression of ALT features, observed upon *PML*/BTR loss, or tolerance of ALT byproducts, observed upon *SLFN11* loss.

## Silencing of *SLFN11* is significantly enriched in *ATRX*-deficient ALT cancers

Our data led us to hypothesise that loss of SLFN11 may co-occur with ALT-positive tumours harbouring *ATRX* loss-of-function mutations. To evaluate this, we conducted immunohistochemistry for SLFN11 protein expression in a cohort of soft tissue tumours (*29*) (including 56 leiomyosarcoma, 36 liposarcoma and 34 osteosarcoma samples) (Fig. 1G). This analysis determined a significant attenuation of SLFN11 in ALT+/*ATRX*-deficient samples (Fig. 1, G and H). Data mining from the cancer dependency (DepMap) project (*30*) also identified *SLFN11* as one of the genes that displayed the strongest transcriptional downregulation in tumour cell lines with damaging or truncating mutations in *ATRX* (fig. S3, A and B). This analysis corroborated previously identified transcriptionally upregulated genes, such as *TSPYL5,* which plays a key role in *ATRX*-deficient/ALT+ models (*31*) (fig. S3A). Furthermore, orthogonal mining of *SLFN11* mRNA expression in cBioportal studies (*32, 33*) revealed it is strongly selected against in sarcoma and glioma tumours with *ATRX* truncating mutations (Fig. 1, I and J). Finally, *SLFN11* mutations are found in ∼8% of uterine corpus endometrial carcinoma (UCEC) and significantly co-occur with *ATRX* mutations (fig. S3, C and D). Together, these data indicate that *SLFN11* loss is significantly enriched across a broad range of *ATRX* deficient cancers, a genetic marker consistently associated with ALT.

## SLFN11 loss permits the emergence of ALT following telomere crisis

Activation of telomere maintenance is essential for cancer cells to escape telomere crisis and acquire unlimited proliferative capacity (*9, 34, 35*). *ATRX* dysfunction is an early event in ALT establishment (*36, 37*), predisposing to telomere crisis escape through ALT initiation via an undefined mechanism (*23, 38*). Since *SLFN11* loss allows tolerance of ALT byproducts and its loss co-occurs with *ATRX* deficient cancers, we sought to determine if *SLFN11* loss influences escape from telomere crisis. To this end, we ablated the catalytic subunit of telomerase, *TERT*, in four genetic backgrounds – WT, *SLFN11*-null, *ATRX*-null and *SLFN11-ATRX*-null – thereby generating compound single (*TERT* KO), double (*SLFN11-TERT* or *ATRX-TERT* DKO) or triple mutants (*SLFN11-ATRX-TERT* TKO, referred to as ‘SAT’ clones) (Fig. 2A and fig. S4, A to C). Six or nine independent clones were selected in *ATRX* proficient or deficient genotypes, respectively. All clones were serially passaged independently in unchallenged conditions for multiple cell divisions until telomere erosion resulted in telomere crisis (Fig. 2B).

**Fig. 2.**
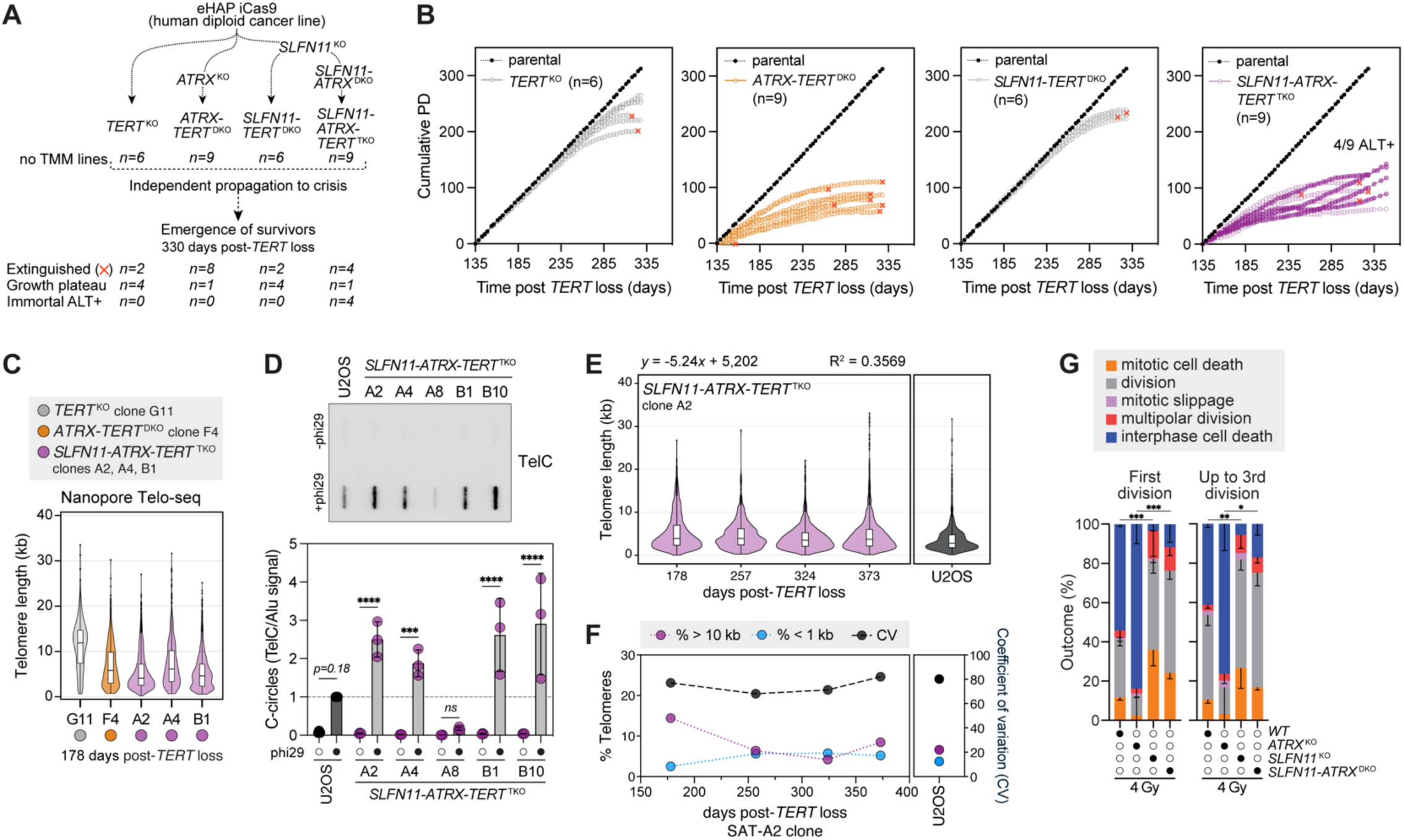
SLFN11 loss permits the emergence of ALT following telomere crisis. **A**, Schematic of *TERT* deficiency clonal evolution experiment. *TERT* ablation resulted in cells with no telomere maintenance mechanism (TMM). Extinguished lines refer to complete cellular extinction; growth plateau to cells still in crisis (have not extinguished and not escaped crisis); immortal refers to cells that activated a TMM to escape crisis. **B**, Growth curves of eHAP cells with deleted *TERT* (parental wild-type *TERT*+ cells included in all graphs). Red cross indicates termination of culture due to complete exhaustion of cellular population. **C**, Violin plot of Telo-seq telomere length measurements at 178 days post-*TERT* loss for clones of the indicated genotypes. Violin represents telomere length distribution. Boxplot shows the median telomere length with IQR and whiskers represent 1.5-fold IQR. Number of telomeric reads per sample and additional details in Supplementary Table 5. **D**, Top: Representative C-circle assay on gDNA from unchallenged asynchronous cultures between 345-377 days post-*TERT* loss. Bottom: Normalised C-circle levels, data are mean ± SD, normalised to U2OS cells (n=3 biological replicates, one-way ANOVA). **E**, Violin plot of Telo-seq telomere length measurements at the indicated timepoint post-*TERT* loss. Violin represents telomere length distribution. Boxplot shows the median telomere length with interquartile range (IQR) and whiskers represent 1.5-fold IQR. Shortening rate equation was calculated by plotting mean telomere length against time, transformed from days to relative PDs using data in Fig. 2B, and performing linear regression analysis. For more details, see Supplementary Table 5. **F**, Telo-seq metrics including telomere length measurements <1 kb or >10 kb or coefficient of variation, at the indicated timepoint post-*TERT* loss. **G**, Extended live cell imaging outcomes of the first (left) and up to third (right) cell division for cells exposed to 4 Gy. Data are mean ± SD (n=3 biological replicates, scoring ≥ 28 cells per replicate, one-way ANOVA with Fisher’s LSD test). Statistics compare mitotic cell death between groups.

Unexpectedly, *ATRX*-null clones entered a crisis-like growth plateau at a much lower population doubling (PD) following *TERT* ablation when compared to *ATRX* proficient cells (Fig. 2B). In all *ATRX*-null clones, entry to telomere crisis occurred independently of SLFN11 and was characterised by reduced proliferative rates, increased telomere length heterogeneity, and stochastic telomere shortening, which was corroborated in clones 178 days post-*TERT* deletion by native long-read sequencing of individual telomeres using Telo-seq (Fig. 2, B and C, and fig. S4, D and E). Relative to *TERT*-null clones, *ATRX*-*TERT*-null clones survived for a longer duration with sub log-phase proliferation before culture extinction or cessation of growth (Fig. 2B). Strikingly, while all *ATRX-TERT*-DKO clones except one were extinguished 330 days after *TERT* loss, 5 out of 9 SAT triple KO clones remained viable, of which four exited crisis and were identified as ALT positive (Fig. 2, B and D). Considering the very low frequency of spontaneous immortalisation through ALT engagement in telomerase deficient cells (0.9 × 10^−8^ to 2.5 × 10^−7^) (*39*), the high rate of ALT+ survivors in SAT clones implicates *SLFN11* loss is a key event required to escape telomere crisis through ALT activation.

Unlike the ALT-primed state that presented muted ALT-like phenotypes (Fig. 1A and fig. S1, D and G), the eHAP-SAT clones that escaped crisis presented robust ALT hallmarks equivalent to well established U2OS ALT+ cancer cells (Fig. 2D and fig. S5, A to D). This includes C-circles, ssTeloC foci, APBs and the presence of ultrabright denaturing FISH telomeric foci suggestive of telomere clustering (*40, 41*). Interestingly, the crisis clone SAT-A8, which did not show robust ALT activity by C-circles or ssTeloC foci, exhibited marked APB levels comparable to the clones that escaped crisis and engaged ALT (Fig. 2D and fig. S5, A to D), suggesting APB formation precedes productive ALT activity.

Using Telo-seq analysis, we corroborated that the telomere length distribution of ALT+ SAT clone A2 resembles that of the U2OS ALT+ cell line (Fig. 2, E and F). Telomeres in *TERT*-null clone G11 shortened ∼46 bp per PD (fig. S5E), comparable to ∼39 bp per PD estimated in IMR90^E6E7^ fibroblasts (*42*). While the telomeres of crisis-state *ATRX-TERT*-null clone F4 exhibited more rapid shortening of ∼ 57 bp per PD; the SAT clone A2, which displayed steady proliferation and no static crisis, had a substantially reduced telomere erosion of ∼5 bp per PD and hallmarks of telomere length heterogeneity (Fig. 2, E and F, and fig. S5E). This is consistent with the SAT-A2 clone engaging in ALT-dependent telomere lengthening to counter natural telomere erosion. Collectively, these data demonstrate that deleting *SLFN11* removes a suppressive barrier key for ALT establishment.

## SLFN11-dependent interphase cell death limits mitotic errors

Critically short telomeres can lose their protective high-order structure and be recognized as chromosome double strand breaks (*43*), triggering mitotic cell death (*44, 45*). To investigate how SLFN11 influences double strand break-induced cell death, we exposed cells to 4 Gy irradiation and performed single-cell analysis using extended live imaging. Irradiation of asynchronous cultures induced interphase lethality prior to mitosis in an SLFN11-dependent manner, which was exacerbated in ALT-primed *ATRX*-null cultures (Fig. 2G, and fig. S6A). Conversely, irradiated *SLFN11*-null cells survived interphase and instead died during the first mitosis or exhibited aberrant divisions (Fig. 2G, and fig. S6, A to C). These data are consistent with SLFN11-dependent interphase death acting as the first line of defence against DNA damage. In the absence of immediate interphase death, *SLFN11* deficient cells accumulated higher levels of multipolar divisions and mitotic slippage, before ultimately dying from mitotic catastrophe (fig. S6, A to C). Our findings indicate that SLFN11 loss does not prevent telomere crisis entry nor DNA damage-induced mitotic cell death, but rather removes an interphase safeguard that eliminates damaged cells before division.

## SLFN11 promotes clearance of *ATRX*-null cells in response to telomeric ssDNA intermediates

Upon treatment with DNA damaging chemotherapeutic agents, SLFN11 promotes replication fork arrest through its ATPase activity (*27, 46*) and induces p53-independent apoptosis via ribosome stalling and ribosome biogenesis impairment, dependent on its tRNA nuclease function (*47, 48*). To determine how SLFN11 imposes the barrier to ALT establishment, we explored the mechanistic basis by which SLFN11 senses, signals and triggers cell death upon ALT priming. To this end, we first assessed the relevance of functional motifs in SLFN11 by mutating the tRNA nuclease catalytic site, the ssDNA-binding motif and the ATPase Walker B motif of SLFN11 (Fig. 3A). *SLFN11-ATRX*-DKO cells were reconstituted with empty vector or mutant forms of SLFN11 and tested for their ability to trigger cell death and ALT priming in response to replication stress (Fig. 3B). *SLFN11-ATRX*-DKO cells expressing Walker B motif mutant E669Q SLFN11 showed elevated cell death comparable to wild-type SLFN11 in *ATRX*-null cells (Fig. 3C). However, *SLFN11* mutations that compromise ssDNA binding (K652D) or tRNA endonuclease activity (E209A) failed to induce cell death in response to replication stress (Fig. 3C). All SLFN11 mutant reconstituted cells exhibited equivalent ssTeloC ALT-priming (fig. S7A). These findings indicate that SLFN11, through its ssDNA binding and tRNA endonuclease activity, but not its ATPase, kills ALT-primed *ATRX*-null cells following replication stress.

**Fig. 3.**
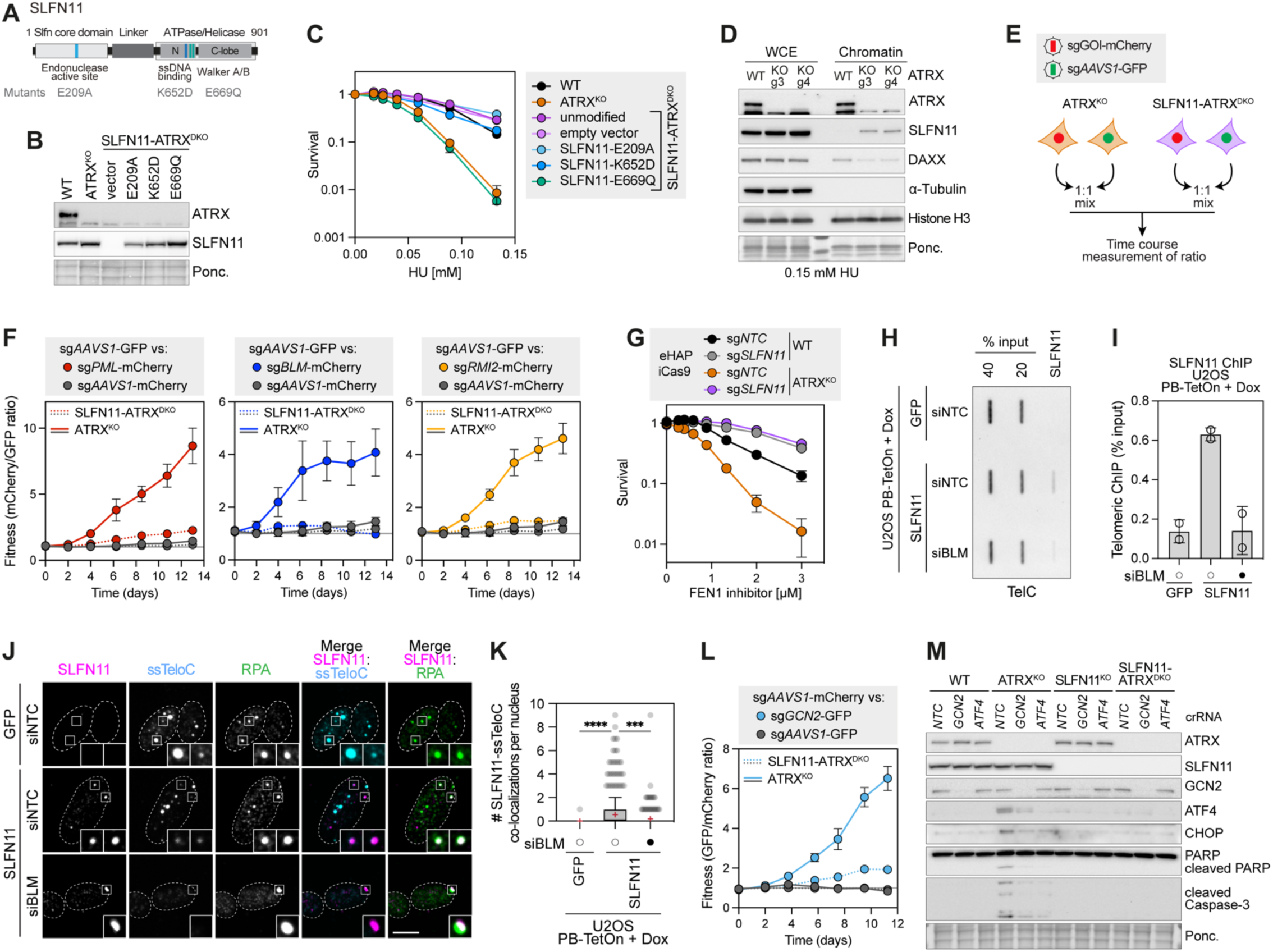
SLFN11 promotes elimination of *ATRX*-null cells in response to telomeric ssDNA intermediates. **A**, Schematic of full-length SLFN11 and mutants used to reconstitute *SLFN11*-*ATRX*-null cells. **B**, Immunoblot of whole cell extracts from cell lines stably expressing *SLFN11* cDNA mutants. Ponceau stain used as loading control. **C**, eHAP cells as in (B) were treated continuously for five days with HU, and viability was determined using CellTiter-Glo. Data are mean ± SEM, normalised to untreated (n=3 biological replicates). **D**, Immunoblot of whole cell extract (WCE) versus chromatin from WT or *ATRX*-null eHAP cells, following 24 h 0.15 mM HU treatment. Alpha-tubulin was used as a soluble fraction loading control. Histone H3 was used as a chromatin fraction loading control. **E**, Schematic of two-colour competitive growth assays. Genes of interest (GOI). **F**, Two-colour competitive growth assays in *ATRX*-null and *SLFN11*-*ATRX*-null cells transduced with the indicated sgRNAs in the presence of Dox. Data for sg*AAVS1*-GFP vs sg*AAVS1*-mCherry are the same in the three graphs. Data are mean ± SEM (n=6 biological replicates). **G**, eHAP cells were treated continuously for five days with FEN1 inhibitor, and viability was determined using CellTiter-Glo. Data are mean ± SEM, normalised to untreated (n=3 biological replicates). **H**, ChIP experiments using antibodies raised against SLFN11. The immunoprecipitated DNA was blotted and probed with a telomere-specific probe. **I**, Quantifications of SLFN11 ChIP signal as % of input at telomeres, in the indicated cell lines as in (H). Data are mean ± SD (n=2 biological replicates). **J**, Representative micrographs for chromatin-bound SLFN11 and RPA immunofluorescence followed by native telomeric PNA FISH. Dashed lines indicate nucleus outlines (as determined using DAPI staining). Scale bar: 10 µm. **K**, Quantification of SLFN11-ssTeloC colocalizations as in (J). Tukey box plots represent individual nuclei, red ‘+’ represents mean (n=2 biological replicates, one-way ANOVA). **L**, Two-colour competitive growth assays in *ATRX*-null and *SLFN11*-*ATRX*-null cells transduced with the indicated sgRNAs in the presence of Dox. Data for sg*AAVS1*-mCherry vs sg*AAVS1*-GFP are the same than in fig. S8F. Data are mean ± SEM (n=6 biological replicates). **M**, Immunoblot of whole cell extracts from unchallenged eHAP cells 3 days after transfection with the indicated crRNAs. Ponceau stain used as loading control.

The requirement for SLFN11 ssDNA binding raised the possibility that the accumulation of ssDNA lesions in *ATRX*-null cells upon ALT priming may lead to SLFN11 recruitment and its activation as a tRNA nuclease. To test this possibility, we examined SLFN11 sub-cellular localisation upon low dose replication stress. HU treatment resulted in chromatin recruitment of both ATRX and its binding partner DAXX (Fig. 3D). Loss of *ATRX* abolished DAXX chromatin recruitment, as expected (*49–51*). Strikingly, in HU-treated *ATRX*-null cells we also observed robust enrichment of SLFN11 on chromatin (Fig. 3D). To test if SLFN11 is activated by lesions that arise during ALT priming, we eliminated ALT activity in *ATRX*-null cells by deleting *PML* or BTR. As shown previously, the cellular fitness of unchallenged *ATRX*-null cells is enhanced upon deleting *PML* or BTR in competitive growth assays (Fig. 1E and Fig. 3, E and F). In contrast, *SLFN11* ablation in *ATRX*-null cells reversed the enhanced cellular fitness observed upon loss of *PML* or BTR (Fig. 3F, and fig. S7B), indicating that in *ATRX*-null cells SLFN11 responds to endogenous lesions generated by PML and BTR complex activities. *ATRX*-null cells were also hypersensitive to chemical inhibition of the lagging-strand DNA flap endonuclease 1 (FEN1) in a SLFN11-dependent manner (Fig. 3G and fig. S7C). Loss of *PML* or BTR desensitised *ATRX*-null cells to FEN1 inhibition (fig. S7C), indicating their activities are responsible for driving the differential SLFN11-dependent lethality observed upon FEN1 inhibition in *ATRX*-null cells. These findings suggest that FEN1-dependent processing protects *ATRX*-null cells from SLFN11 activation, whereas excessive strand displacements by BTR within PML bodies trigger SLFN11 activity in *ATRX* deficient cells.

We further investigated whether SLFN11 directly localises at telomeric ssDNA, which is a unique hallmark of ALT+ cells (*14*). To test this possibility, we inducibly re-expressed SLFN11 in ALT+ *ATRX-SLFN11*-deficient U2OS cells (fig. S7, D and E). Anti-SLFN11 monoclonal antibodies were used to chromatin immunoprecipitate (ChIP) SLFN11 protein in inducible U2OS cells. SLFN11 interacted with telomeric DNA in a BLM-dependent fashion (Fig. 3, H and I, and fig. S7, F to H). As an orthogonal approach, we tested whether SLFN11 accumulates at telomeric ssDNA foci by co-staining for SLFN11, RPA and ssTeloC. We observed BLM-dependent co-localisation of SLFN11 with a subset of telomeric ssDNA foci that were also positive for RPA (Fig. 3, J and K, fig. S7I). Collectively, these data establish that SLFN11 senses telomeric ssDNA lesions that arise in *ATRX* deficient cells during ALT.

## SLFN11-ISR axis is responsible for *ATRX*-null cell death upon endogenous damage

Next, we explored the role of the SLFN11 tRNA endonuclease activity, which is also required to induce cell death in *ATRX*-null cells upon ALT priming (Fig. 3C). A recent study has shown that SLFN11 cleaves rare tRNA^Leu^-UUA in response to certain chemotherapeutics, thereby inducing ribosome stalling and triggering a global decrease in protein synthesis that precedes apoptosis (*47*). To determine if a comparable mechanism occurs in *ATRX*-null cells upon ALT priming, we conducted nascent peptide labelling with puromycin, coupled with single-cell flow cytometry analysis. This revealed a significant fraction of cells undergoing translation inhibition in *ATRX*-null cells within 12 to 16 hours upon ALT priming, which correlated with high levels of cleaved-caspase 3 (fig. S8, A to C). By harnessing ribosome profiling in conjunction with differential ribosome measurements of codon reading (diricore) (*52*), we observed that ALT priming in *ATRX*-null cells leads to pronounced ribosome stalling at Leu^UUA^ codons in a SLFN11-dependent manner (fig. S8D), similarly to that seen with treatment with etoposide (*47*).

While the SLFN11 response to chemotherapeutics activates ZAKα and induces ribotoxic stress response markers including phosphorylation of MAP2K4/7 and JNK (*47, 48*), no such induction was observed in *ATRX*-null cells upon ALT priming (fig. S8E). This suggested a distinct signalling mechanism promoting cell death in *ATRX*-null cells upon ALT priming. Consistent with this possibility, the GCN2/*EIF2AK4* kinase and its partner GCN1/*GCN1L1* were amongst the top enriched hits in our screens, which are known to sense ribosome stalling as part of the integrated stress response (ISR; Fig. 1, C and D) but are dispensable for the induction of cell death in response to etoposide (*47*). To explore a potential role for the ISR in signalling cell death during ALT priming, we asked whether *SLFN11* loss is epistatic with *GCN2* and *GCN1* loss in promoting cell death in *ATRX*-null cells. Using two-colour competitive growth assays, genetic loss of *GCN2* or *GCN1* conferred increased cellular fitness in unchallenged *ATRX*-null cells, but not in the *SLFN11-ATRX*-DKO cells consistent with epistasis (Fig. 3L, and fig. S8, F and G). Similar to *PML* or BTR, loss of *GCN2* or the downstream target of ISR *ATF4*, desensitised *ATRX*-null cells to FEN1 inhibition (fig. S8H).

In agreement with the involvement of *GCN2* and *GCN1,* we observed accumulation of downstream markers of the ISR including ATF4 or CHOP associated with apoptosis in a SLFN11-dependent manner (Fig. 3M). Hence, unlike acute high DNA damage conditions where the ribotoxic stress response mediates apoptosis (*47, 48*), death of ALT-primed *ATRX*-null cells driven either by endogenous damage or inhibition of FEN1 is mediated by prolonged activation of the ISR (Fig. 3M and fig. S8H). Collectively, these results demonstrate that upon ALT priming, SLFN11 is activated as a tRNA nuclease resulting in ribosome stalling at rare leucine-encoding UUA codons, this activates translation inhibition by *GCN1/GCN2* and triggers apoptosis via ATF4.

## SLFN11 loss in *ATRX*-null tumours confers naive therapy-resistance

ALT cancers are typically aggressive and refractory to current treatments (*53*). Small molecule inhibitors of the replication checkpoint kinase ATR have been suggested to kill *ATRX* deficient ALT+ cells, but its application is controversial (*54–57*). Having established *SLFN11* loss as a frequent occurrence in ALT cancers, we hypothesised that the genetic events needed to drive ALT could be responsible for tumours being innately resistant to targeted therapies. To formally test whether *SLFN11* loss renders tumours resistant to low-dose ATR inhibitor AZD6738 administered at a continuous daily schedule *in vivo*, we developed isogenic eHAP WT and ALT-primed *ATRX*-null cells expressing sg*NTC* or sg*SLFN11* and iCas9 as subcutaneous xenografts in immunodeficient mice (Fig. 4A). As the tumours developed, mice were treated with doxycycline to induce *SLFN11* loss and once a median tumour size of 100 mm^3^ was reached, mice were randomised and treated with 25 mg/kg AZD6738 until the endpoint. The growth of *ATRX*-null tumours was significantly inhibited by single-agent AZD6738, while the inhibitor showed no activity in *ATRX* proficient tumours (Fig. 4, B and C, and fig. S9, A to C). Strikingly, *SLFN11* loss in *ATRX*-null tumours conferred ATR inhibitor resistance preventing tumour stasis or regression (Fig. 4, B and C, and fig. S9, A and B). Notably, *ATRX-SLFN11*-null tumours accumulated the highest levels of DNA damage upon ATR inhibition, despite their tumour progression being indistinguishable from *ATRX*-proficient counterparts and vehicle controls (Fig. 4, D and E). This demonstrates that *SLFN11* loss permits the evolution of *ATRX* deficient tumours that are innately resistant to ATR inhibitors and suggests this may be the case for other targeted therapies.

**Fig. 4.**
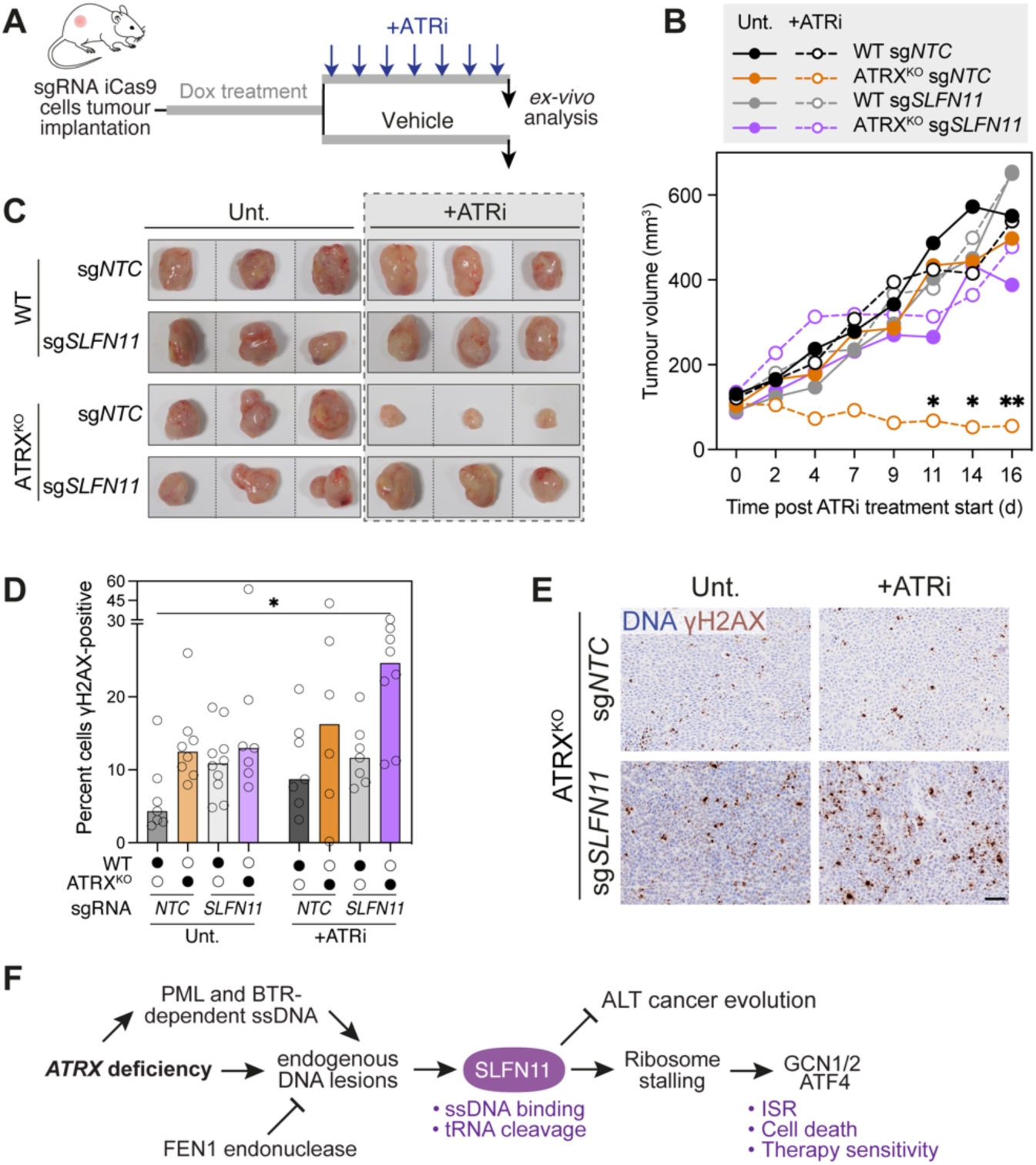
SLFN11 loss in *ATRX*-null tumours confers naive therapy-resistance. **A**, Schematic of tumour xenograft experiment. **B**, Tumour growth of eHAP iCas9 sgRNA xenografts either untreated or treated with 25 mg/kg AZD6738. Data are median (n=6-10 animals per group). Statistics compare ATR inhibitor treated *ATRX*-null tumours (sg*NTC* vs sg*SLFN11*) using multiple unpaired t tests. **C**, Representative pictures of tumours analysed ex-vivo at the experiment endpoint as in (A-B). **D**, Quantification of the percent of cells positive for γH2AX in a tumour section. Data are scatter dot plot of individual tumours, with bar at median (n=6-10 tumours per group, one-way ANOVA). **E**, Representative images of γH2AX immunohistochemistry on xenografts. Scale bar: 50 µm. **F**, Schematic model of how SLFN11 acts as a barrier to eliminate pre-ALT *ATRX*-deficient cells with endogenous stress. Its loss results in ALT tumour evolution, replication stress tolerance and innate therapy-resistance.

## Discussion

The DNA damage-activated tRNA nuclease SLFN11 is epigenetically silenced in about half of all cancers and is the strongest predictor of chemoresistance. Why cancer cells frequently inactivate SLFN11 is unknown. Here, we demonstrate that SLFN11 responds to endogenous telomere replication stress and serves as a barrier to the acquisition of ALT, triggering p53-independent apoptosis (Fig. 4F). During clonal evolution, we define that absence of *TERT*, *ATRX* and *SLFN11* constitute the minimal genetic requirements for ALT establishment. The frequent occurrence of SLFN11 loss in *ATRX*-null ALT cancers suggests that the same process occurs during cancer evolution in patients. Our work provides a framework that reconciles the use of ATR inhibitors to target *ATRX* deficiency and ALT (*54–58*), where SLFN11 status is a determinant of ATR inhibitor therapy response. Our findings show that *SLFN11* loss overcomes the challenge imposed by heightened toxicity of PML-BLM dependent telomere recombination intermediates, allowing ALT establishment at the expense of genome integrity and innate chemoresistance.

Our results identified *ATRX* loss as a determinant of rapid telomere shortening resulting in a decreased time to reach telomere crisis and increased telomere heterogeneity. It is reasonable to suggest that telomere shortening may reflect the stochastic occurrence of ssDNA at telomeres. During telomeric DNA replication, most telomeres are copied by a single replication fork, initiated primarily from within the subtelomeric region (*59*). As such, potential defects in lagging strand synthesis in *ATRX*-null cells, such as un-ligated Okazaki fragments (*17, 26*), would present as gapped leading strand templates during the next replication cycle. This could result in one-ended DNA breaks, stopping telomere replication prematurely and possibly leading to replisome loss, which would stochastically shorten the telomere.

Notably, *TP53* and *RB1* are two tumour suppressors often mutated in ALT cancers, which favour or enable ALT activation by overcoming the cellular senescence barrier, which occurs earlier than the crisis escape barrier (*25, 60–62*). SLFN11’s uniqueness lies in the fact that a single protein harnesses a specific ssDNA sensing module, rapidly translocates to the cytoplasm upon ssDNA recognition (*63*), and executes cell death through cleavage of tRNA^Leu^(UUA), depletion of which inflicts ribosome stalling and activation of the downstream ISR. In ALT-primed *ATRX*-null cells, we show inhibition of FEN1-dependent processing of flap structures, possibly in Okazaki fragments, triggers SLFN11/GCN2/ATF4-dependent cell death. In the absence of exogenous replicative stress, spontaneous engagement of telomeres with PML and the BTR complex in ALT-primed cells confers cytotoxicity due to activation of SLFN11. Similarly, we demonstrate that acquisition of ALT as a mechanism to escape critical telomere erosion is restricted by SLFN11 activity, and exogenous re-expression of SLFN11 engages with ALT-specific telomeric ssDNA species in ALT+/*ATRX*-*SLFN11*-deficient U2OS cells. SLFN11 has also been proposed as an immunostimulatory ssDNA pattern recognition receptor (*63*), but whether this function is important in the context of ALT is unclear. Notably, the cytosolic double-stranded DNA sensor cGAS and the downstream adaptor protein STING have been shown to mediate type I interferon responses in ALT+ cells, triggered by the presence of cytosolic telomeric DNA fragments (*64, 65*). In addition to inducing cell death, the binding of SLFN11 to telomeric ssDNA intermediates could directly restrict recombination in pre-ALT cells by limiting the binding of RAD51/RAD52. Such a role is consistent with observations that SLFN11 suppresses sister chromatid exchanges and inhibits homologous recombination in a gene conversion assay (*46*).

The barrier imposed by SLFN11 to ALT establishment and tumour evolution in the presence of telomeric replicative stress appears to resemble the impact of DNA damage checkpoints in the oncogene-induced DNA damage model of cancer development (*66, 67*). In both instances, key tumour suppressor pathways must be lost to allow continued proliferation in the presence of endogenous genome damage. *SLFN11* expression is sufficient to predict response to chemotherapeutics in cancer (*2, 5*), likely though direct induction of p53-independent apoptosis in response to exogenous replication stress conferred by cytotoxic chemotherapeutics. By extrapolation beyond ALT, oncogene activation or mutations in key tumour drivers associated with heightened replication stress may also select for loss of *SLFN11* early in tumour evolution, thus explaining the widespread silencing of *SLFN11* across many cancers. Indeed, *SLFN11* loss is significantly associated with cancer cell lines harbouring *ARID1A, SMARCA4* or *MSH6* mutations (fig. S9D), suggesting that for instance microsatellite instability could select against *SLFN11* expression. We predict this comes at a cost of both increased chromosome segregation errors as damaged cells are not removed in interphase and enhanced naive chemoresistance. Our findings identify endogenous replication stress as the reason for the frequent *SLFN11* inactivation in cancer, which provides an explanation for why ALT-positive tumours are generally refractory to chemotherapy and have a poor prognosis.

## Acknowledgments

We thank members of the Boulton lab, J.B. Vannier, T.H. Stracker, S.C. West and C. Swanton for critical reading of the manuscript and the Francis Crick Institute’s Genomics, Advanced Light Microscopy, Cell Science, Flow Cytometry, Bioinformatics & Biostatistics, Experimental Histopathology and High Throughput Screening science technology platforms for support. We also thank AstraZeneca and Artios Pharma for providing reagents (AZD6738 and ART3999, respectively). S.S-B. was supported by an EMBO Long Term Fellowship (ALTF 707-2019) and a MSCA individual fellowship (886577). G.H. is supported by the Radiation Research Unit at the Cancer Research UK City of London Centre Award (C7893/A28990). Work from the A.J.C. lab is supported by the Australian Research Council (FT210100858, DP240101869) and enabled by the ACRF Telomere Analysis Center. Work in the S.J.B. lab is supported by the Francis Crick Institute (CC2098), European Research Council Advanced Investigator grants (TelMetab, ChrEndProt), a Wellcome Trust Senior Investigator Award, and CRUK RadNet City of London.

## Materials and Methods

### Cell lines

eHAP cells (Horizon Discovery, stable diploid cells used throughout) and HT1080 cells (ATCC) were modified by a lentiviral integration of doxycycline-inducible Cas9 nuclease (iCas9) with an Edit-R inducible lentiviral Cas9 vector (Horizon Discovery)(*^68^*). iCas9 cell lines were derived as single clones from the pool of transduced cells after blasticidin selection and were chosen based on a high cutting efficiency and negligible leakiness using a BFP/GFP reporter assay (Addgene #67980). eHAP cell lines were grown in Iscove′s modified Dulbecco′s medium (IMDM) with 10% Tet-free FBS and 1% penicillin/streptomycin. HT1080 cell lines were grown in Eagle’s minimum essential medium (EMEM) with 10% Tet-free FBS and 1% penicillin/streptomycin. The sarcoma cell line U2OS (ATCC) was grown in McCoy’s 5A medium with 10% FBS and 1% penicillin/streptomycin. All cell lines were grown at 37°C in 5% CO_2_. For clonal evolution experiments, clonal representation was maintained at 5×10^4^ to 10^5^ cells per clone during passaging. The number of population doublings (PD) was calculated at each passage as PD = 3.32(log [number of counted cells/number of cells seeded]).

### Reagents and Antibodies

Reagents used in this study were DMSO (Sigma), hydroxyurea (Sigma), aphidicolin (Sigma), ATR inhibitor AZD6738 (Selleckchem), FEN1 inhibitor ART3999 (Artios Pharma), anisomycin (Sigma, 5 μg/ml) or doxycycline (Sigma, 1 μg/ml). Antibodies used in this study were: anti-alpha-Tubulin (Sigma T6074), anti-ATF4 (Cell Signaling Technology 11815), anti-ATRX (Bethyl Laboratories A301-045A), anti-BLM (Abcam ab2179), anti-Cleaved Caspase 3 D175 (for immunoblot, Cell Signaling Technology 9664), anti-Cleaved Caspase 3 (for Flow, BD Biosciences 559565), anti-CHK1 (Cell Signaling Technology 2360), anti-p-CHK1 S345 (Cell Signaling Technology 2348), anti-CHOP (Cell Signaling Technology 2895), anti-DAXX (Santa Cruz sc-7152), anti-GCN1L1/GCN1 (Bethyl Laboratories A301-843A), anti-GCN2 (Cell Signaling Technology 3302), anti-HA (Roche 11867423001), anti-Histone H3 (Abcam ab10799), anti-MCL1 (Cell Signaling Technology 5453), anti-p-JNK (Cell Signaling Technology 4668), anti-p-MAP2K4 (Cell Signaling Technology 4514), anti-p-p53 S15 (Cell Signaling Technology 9284), anti-PARP (Cell Signaling Technology 9542), anti-PML (for immunoblot, Bethyl Laboratories A301-167A), anti-PML (for immunofluorescence, Santa Cruz sc-966), anti-Puromycin (Merck MABE343), anti-RMI2 (Abcam ab122685), anti-RPA (Abcam ab2175), anti-SLFN11 (for immunohistochemistry, IF and ChIP, Cell Signaling Technology 34858), anti-SLFN11 (for immunoblot, Santa Cruz sc-374339), anti-Vinculin (Abcam ab11194), anti-Digoxigenin-AP Fab fragments (Roche 11093274910), Goat polyclonal anti-mouse Immunoglobulins/HRP (Agilent-Dako P0447), Swine polyclonal anti-rabbit Immunoglobulins/HRP (Agilent-Dako P0399), Goat anti-mouse IgG (H+L) Alexa Fluor 488 (Thermo Fisher Scientific A11001), Goat anti-rabbit IgG (H+L) Alexa Fluor 647 (Thermo Fisher Scientific A21245).

### Plasmids

cDNA for *ATRX*-HA and *SLFN11* were introduced into pDONR221 (Invitrogen) by amplification using attb1/attb2 primers (Supplementary Table 1). Mutant variants were introduced by Q5 site directed mutagenesis (NEB) in pDONR221, according to manufacturer’s instructions, using primers in Supplementary Table 1. Mammalian expression vectors for *ATRX*-HA or untagged *SLFN11* variants were made in a piggyBac (PB) transposon delivery system PB-EF1α-DEST-IRES-Puro for stable expression or PB-TetOn-DEST-IRES-Puro for Dox-inducible expression. cDNAs were cloned into PB vectors using the Gateway technology (Thermo Fisher Scientific) according to the manufacturer’s protocol. Expression constructs were introduced into eHAP or U2OS cells by co-transfection with Super PiggyBac Transposase (PB200A-1, System Biosciences).

### Generation of CRISPR full knockout cell lines

Isogenic knockouts of *ATRX* in eHAP iCas9 cells were made by transfection with synthetic tracrRNA/crRNA (Horizon Discovery). Upon transfection cells were incubated with 1 µg/mL doxycycline to induce Cas9 expression for 72 hours, then cells were seeded by limiting dilution in 96-well plates to derive as single cell clones. Isogenic wild-type/*ATRX* knockout in HT1080 iCas9 cells were made by transducing with *NTC* or *ATRX* sgRNAs cloned into lenti-sgRNA-hygro (Addgene #104991). Cells were selected in hygromycin, followed by treatment with 1 µg/mL doxycycline to induce Cas9 expression for 72 hours and then seeded by limiting dilution in 96-well plates to derive as single cell clones. Isogenic knockouts of *SLFN11* and *TERT* in eHAP iCas9 cells were made by transient transfection with lenti-sgRNA-puro (Addgene #104990) or lenti-sgRNA-hygro (Addgene #104991), respectively. Upon transfection cells were incubated with 1 µg/mL doxycycline to induce Cas9 expression for 72 hours and selected for 48 hours in puromycin (0.4 µg/mL) or hygromycin (0.4 mg/ml) after 24 hours of transfection. Then cells were seeded by limiting dilution in 96-well plates to derive as single cell clones. All clones were validated by immunoblotting (except TERT where antibodies were unavailable) and genomic DNA sequencing. Targeting sgRNA sequences are listed in Supplementary Table 2.

### Lentiviral transductions

To produce lentivirus, 900,000 293FT cells were seeded in a 6-well plate and then transfected with packaging plasmids (566 ng of pLP1, 266 ng of pLP2, 370 ng of pLP/VSVG) along with 1 µg of lentiviral vector plasmid using 4 µL Lipofectamine 2000 (Thermo Fisher Scientific) as per the manufacturer’s instructions. Medium was refreshed 18 hours later. Virus-containing supernatant was collected 72 hours post transfection, cleared through a 0.45-µm filter, supplemented with 8 µg/ml polybrene (Sigma), and used for infection of target cells.

Transductants were selected after one day of recovery. The following antibiotics were used for selection of transductants: puromycin (eHAP 0.4 µg/mL; HT1080 1 µg/mL; 2-3 days), hygromycin (eHAP 0.4 mg/ml, HT1080 0.1 mg/mL; 5 days) and blasticidin (eHAP 8 µg/mL, HT1080 5 µg/mL; 5 days).

### Generation of doxycycline-inducible Cas9 knockout cell lines

Inducible CRISPR knockout cell lines were generated by transducing iCas9 cells with lentivirus produced from the lenti-sgRNA-puro construct (Addgene #104990), target sequences of sgRNAs listed in Supplementary Table 2. Inducible knockout of target proteins was confirmed in the pooled cell line by immunoblotting following treatment with 1 µg/mL doxycycline for 96 hours.

### Native FISH (ssTeloC)

Cells were grown on 96-well microplates (PerkinElmer) and fixed in 4% formaldehyde for 10 min at room temperature (RT). After fixation, cells were washed with PBS twice and processed for native FISH. Cells were incubated with 250 μg/mL RNaseA in ADB blocking solution (Antibody Dilution Buffer; 10% goat serum, 0.1% Triton X-100, 0.1% saponin in PBS) for 1 h at 37°C. Next, plates were dehydrated in 70%, 85%, and 100% (v/v) ethanol for 5 minutes each and then air-dried. Plates were hybridized with a telomeric TelG-TAMRA PNA probe (PNA Bio) in hybridizing solution (70% formamide, 1 mg/mL blocking reagent (Roche), 10 mM Tris-HCl pH 7.5) for 2 hours at RT and washed twice with 2X SSC with incubation for 10 min with DAPI in the last wash, followed by two washes with PBS. Plates were sealed and scanned using an Opera Phenix Plus High-Content Screening System (PerkinElmer), where images were captured using 40x objective and analysed using Harmony High-Content Imaging and Analysis Software (PerkinElmer).

### C-circle assay

The C-circle assay protocol was adapted from Henson et al(*69*), by using the quick C-circle preparation (QCP) protocol. DNA concentration was measured by fluorimetry using the Qubit dsDNA HS Assay (Thermo Fisher Scientific). Rolling circle amplification by phi29 DNA Polymerase (Thermo Fisher Scientific) was performed by incubating 30 ng of DNA in a thermocycler at 30°C for 8 hours, after which polymerase was inactivated at 70°C for 20 min. For slot blot detection, samples were blotted onto Hybond N+ positively charged nylon membranes (Amersham) under native conditions. After crosslinking, membranes were hybridized with either with non-radioactive 3’ DIG labelled probes, as previously described(*70*), or with radioactive labelled probes, as previously described(*65*). For chemiluminescence acquisition we used a ChemiDoc MP imaging system (Bio-Rad). Membranes hybridised with γ-^32^P-ATP labelled probes were exposed onto a phosphorimaging plate (GE Healthcare) and scanned using Typhoon FLA 9500 (GE Healthcare).

### CRISPR/Cas9 screening

eHAP iCas9 wild-type or ATRX^KO^ cells were transduced in three biologically independent transductions at a multiplicity of infection (MOI) of 0.4 with the lentiviral Brunello library (Addgene #73179-LV, sgRNA only vector), with a theoretical coverage of 500 cells per sgRNA. Transduced cells were selected with puromycin (0.4 µg/mL) for 48 hours. Cells were subcultured every two days in doxycycline (1 µg/mL) for the initial 6 days and later either subcultured every two days without doxycycline (unchallenged) or in 0.06 mM HU, while keeping the theoretical library coverage of 500 cells per sgRNA. Cells were collected 16 days after doxycycline addition (for HU treated cells that is 10 days after HU addition), when cell pellet samples of 60 million cells were stored at –80°C. Genomic DNA was isolated with the PureLink Genomic DNA Mini Kit (Thermo Fisher Scientific). 200 µg of genomic DNA was then used for library preparation, with one-step amplification of genome-integrated sgRNAs by using P5 mix and P7 barcoded oligonucleotides in a PCR reaction with Ex Taq polymerase (TaKaRa) (Supplementary Table 1). PCR products were purified by agarose gel extraction using the QIAquick Gel Extraction Kit (Qiagen) followed by the MinElute PCR Purification Kit (Qiagen). Libraries were sequenced using Illumina HiSeq 4000 with 100 bp reads (30 million reads per sample).

### CRISPR sequencing analysis

Raw data was trimmed by obtaining 20 bp after the first occurrence of ‘‘CACCG’’ in the read sequence. Trimmed reads were then mapped with BWA (version 0.5.9-r16)(*71*) to a database of guide sequences for the human CRISPR Brunello lentiviral pooled library downloaded from Addgene (https://www.addgene.org/pooled-library/broadgpp-human-knockout-brunello/) with the parameters ‘‘-l 20 –k 2 –n 2’’. sgRNA counts were obtained after filtering the mapped reads for those that had zero mismatches and mapped to the forward strand of the guide sequence. The MAGeCK ‘test’ command (version 0.5.7)(*72*) was used to perform the sgRNA ranking analysis between the relevant conditions with parameters ‘‘–norm-method total–remove-zero both’’.

### Two-colour competitive growth assays

eHAP iCas9 cells were transduced with virus particles expressing sgRNAs either in lenti-sgRNA-GFP-NLS-P2A-puro or lenti-sgRNA-mCherry-NLS-P2A-puro constructs, target sequences of sgRNAs listed in Supplementary Table 1. Selected transductants were mixed 1:1 (15,000 cells each) and seeded in a 24-well plate. Cells were imaged for GFP and mCherry signals in an Incucyte S5 System (Sartorius). Cells were subcultured every 3-4 days when near-confluency was reached. The “Basic Analyzer” fluorescent object ratio analysis tool was used to tailor segmentation and quantification. 16 images were taken per timepoint per well per replicate.

### Immunohistochemistry (IHC) of Tissue microarrays (TMA)

3µm FFPE TMA sections of soft tissue tumours were baked for 1 hour at 60°C before IHC staining performed using Leica Bond Rx autostainer platform. Sections were stained with SLFN11 antibody (Cell Signaling Technology, 34858, clone D8W1B) at 1:200 dilution for 15 minutes at RT. Epitope antigen retrieval solution 2 (Leica, AR9640) was used for 20 min at 95°C prior the antibody application. BOND Polymer Refine Detection kit (Leica, DS9800) was used, including hydrogen peroxidase, anti-rabbit polymer, DAB and haematoxylin counterstain. Slides were coverslipped using Tissue-Tek GlasTM g2 Automated Glass Coverslipper. Stained TMA sections were scanned using the Olympus VS200 ASW whole slide scanner at 20x magnification. The scanned images were de-arrayed within QuPath and all cores examined during the scoring process to exclude manually those with either no tumour represented or with artefacts (tissue folding, for example) precluding assessment. Cells were detected using the QuPath cell detection module and numbers of SLFN11 positive nuclei per TMA core were quantified using DAB intensity measurement in the identified cells.

### DepMap analysis

RNAseq read count data from RSEM (unstranded mode) (OmicsExpressionGenesExpectedCountProfile.csv) were downloaded from DepMap release 22Q4 and normalised across cell lines using a variance stabilizing transformation (VST). Samples were stratified by *ATRX* mutation status, derived from the somatic point mutations and indels called in the DepMap cell lines (OmicsSomaticMutations.csv, “VariantInfo” column). Cell lines were considered *ATRX*^mut^ when carrying damaging/truncating mutations classified by FRAME_SHIFT_DEL, FRAME_SHIFT_INS or NONSENSE. Cell lines were considered wild-type when not carrying an *ATRX* mutation. Cell lines carrying *ATRX* variants of unknown significance such as missense, silent, splice site or in-frame deletions/insertions were excluded. We investigated significant changes in gene expression among cellular models using a Wilcoxon rank sum test to compare VST values between groups (Supplementary Table 4). Plots were generated using R version 4.2.3 and ggplot2.

A fast Gene Set Enrichment Analysis (fGSEA) approach was used to identify candidate genes with a damaging mutation status associated with *SLFN11* expression in each tissue lineage (https://github.com/ctlab/fgsea). All data were downloaded from DepMap release 24Q4. RNAseq read count data from RSEM unstranded mode (OmicsExpressionGenesExpectedCountProfile.csv) were rounded to integers and stratified by DepMap cell line tissue of origin according to the OncotreeLineage column of the available metadata (Model.csv). Each tissue specific counts matrix was then normalised using DESeq2’s variance stabilizing transformation (VST). The mutational status of each gene in each of the DepMap tissue cell lines was determined. “Damaging” mutations were assigned based on a score >0 in the DepMap OmicsSomaticMutationsMatrixDamaging.csv table. If a cell line did not carry a damaging mutation for a gene but did harbour another somatic point mutation or indel as defined in the VariantInfo columns of the DepMap OmicsSomaticMutations.csv table, it was defined as “other”. Genes that were free of any mutation to the query gene were assigned a mutational status of “none”.

fGSEA was used to look for a significant association between SLFN11 expression and the damaging mutational status of cell lines for each query gene. The input for fGSEA was (i) vector of normalised RNA-seq counts for SLFN11 across tissue-specific DepMap cell lines, mean-centered and (ii) a list of tissue-specific cell lines carrying a damaging mutation for a given query gene. Note that only cell lines of mutational status “damaging” or “none” were included in the ranked list to avoid mutation types oof unknown function. A minimum threshold of >=5 cell lines carrying damaging mutations was imposed prior to testing. fGSEA calculates an enrichment Score (ES) indicative of the degree to which the cell lines carrying a damaging mutation for a particular query gene is enriched or overrepresented at either the top or bottom of the ranked list of SLFN11 expressing cell lines. Mutational labels were subjected to 1000 permutations in order to generate a null distribution of ES values and enable p-value calculation. The process was repeated for each gene within each tissue subset. Adjusted p-values were calculated per tissue using the Benjamini-Hochberg (BH) procedure.

### Clonogenic survival assay

400 eHAP iCas9 cells per well were seeded in 24-well plates in technical triplicate, colonies were grown for 6 days. 1000 U2OS cells per well were seeded in 6-well plates in technical duplicate, colonies were grown for 13 days. Colonies were then fixed and stained with 0.5% crystal violet solution with 20% methanol. Plates were scanned and analysed using GelCount (Oxford Optronics).

### Whole cell extracts

Cells were rinsed with PBS, trypsinised and collected in growth medium. Cells were pelleted by centrifugation at 500 g for 5 min and washed once with PBS. Cell pellets were frozen on dry ice and stored at –80°C. For lysis, cell pellets were thawed on ice and resuspended in RIPA Buffer (10 mM Tris-Cl pH 8.0, 1 mM EDTA, 0.5 mM EGTA, 1% Triton X-100, 0.1% sodium deoxycholate, 0.1% SDS, 140 mM NaCl, 1X phosphatase (PhosSTOP, Roche) and protease (cOmplete, EDTA-free, Roche) inhibitor mixes) and incubated on ice for 20 min. Lysates were sonicated with a probe at medium intensity for 10 seconds in a Soniprep 150 instrument and clarified by centrifugation at 13,000 g for 15 min at 4°C. Protein concentration was determined using the DC Protein Assay (Bio-Rad) according to the manufacturer’s instructions. Proteins were denatured in 2X NuPAGE LDS sample buffer (Invitrogen) and 1% 2-mercaptoethanol (Sigma) for 5 min at 95°C.

### SDS-PAGE and immunoblotting

Proteins were separated by SDS-PAGE using NuPAGE mini gels (Invitrogen) and transferred onto 0.2 µm pore Nitrocellulose membrane (Amersham Protran; Sigma) using standard procedures. Membranes were blocked with 5% skim milk/TBST (TBS/0.1% Tween-20) for 1 hour at RT and probed with the indicated primary antibodies overnight at 4°C. Membranes were then washed 3 times with TBST, incubated with appropriate secondary antibodies conjugated to a horseradish peroxidase (HRP) for 1 hour at RT and washed again 3 times with TBST. Immunoblots were developed using Clarity or Clarity Max Western ECL Substrate (Bio-Rad). Chemiluminescence was acquired using a ChemiDoc MP imaging system (Bio-Rad).

### Chromatin fractionation

Cells seeded 36-48 hours prior to collection were treated with 0.15 mM HU for 24 hours. Cells were trypsinised and resuspended in ice-cold PBS. 3 million cells were kept on ice for whole cell lysate control. 3 million cells were spun down for 5 min at 500 g, and resuspended in 200 µL CSK buffer (10 mM PIPES pH 7.0, 100 mM NaCl, 300 mM sucrose, 1.5 mM MgCl_2_, 5 mM EDTA, 0.5% Triton X-100, 1X phosphatase (PhosSTOP, Roche) and protease (cOmplete, EDTA-free, Roche) inhibitor mixes) and incubated on ice for 10 min. Cells were spun down at full speed for 10 seconds and the supernatant (soluble fraction) was collected. The chromatin pellet was washed in 500 µL of CSK buffer. Chromatin pellets and whole cell pellets were resuspended in 150 µL 1X NuPAGE LDS sample buffer (Invitrogen) and 1% 2-mercaptoethanol (Sigma). All samples were boiled at 95°C for 10 min and sonicated with a probe at medium intensity for 10 seconds in a Soniprep 150 instrument.

### Global translation and apoptosis analysis by flow cytometry

To visualize global translation, puromycin incorporation was performed as described(*47*) by treating cells with medium containing puromycin (2 μg/ml) for 10 minutes before washing and harvesting cells. Cells were fixed in Fix buffer I (BD Biosciences) for 10 minutes at 37°C, subsequently permeabilized with Perm Buffer III (BD Biosciences) on ice for 30 minutes and blocked in PBS with 10% goat serum for 30 minutes at RT. Cells were incubated with primary antibodies in blocking buffer for 1 hour at RT followed by incubation with Alexa Fluor conjugated secondary fluorescent antibodies along with DAPI in blocking buffer for 1 hour at RT. Cells were analysed in an LSRFortessa Cell Analyser (BD Biosciences). As previously described for puromycin staining(*47*), the laser intensity was slightly adjusted between measurements. Samples were analysed using FlowJo software (BD Biosciences).

### Ribosome profiling

The ribosome-protected fragments libraries were prepared as previously described (*73*), without harringtonine treatment. Trizol (Invitrogen) was used to isolate total RNA, following manufacturer’s instructions. Quick gel extraction of RNA from polyacrylamide gels was performed followed by overnight extraction at –80°C. Depletion of rRNA was not performed and rRNA was afterwards removed during data processing. The libraries were analysed on a 2100 Bioanalyzer using a 7500 chip (Agilent, Santa Clara, CA), quantified using the KAPA qPCR Library Quantification Kit (Roche, KK4824) and sequenced with 75-base single-reads using a High Output Kit v2.5 (75 cycles) (Illumina, Inc.).

### Live cell imaging and analysis

Differential interference contrast (DIC) microscopy was performed on a Zeiss Cell Observer wide field microscope, with a 20x 0.8 numerical aperture air objective and Axiocam 506 monochromatic camera using ZEN Blue v2.6 (Zeiss), at 37°C, 10% CO_2_ and 3% O_2_. Before imaging, cells were irradiated using an X-RAD 320 (1 Gy min^-1^; Precision X-Ray) irradiator at Westmead Institute of Medical Research (WIMR). All imaging was performed in the CMRI Telomere Analysis Center (ATAC), and images were acquired every 6 min for a total duration of 72 hours. Single cell outcomes were assigned based on morphological features and mitotic duration was scored from nuclear envelope breakdown until cytokinesis or mitotic cell death using ZEN Blue v2 or v2.6 (Zeiss).

### Sensitivity to DNA damaging agents

200 cells per well were seeded in white opaque 96-well plates (Greiner). Cells were treated the following day with the range of concentrations indicated for each compound. After 5 days of treatment, survival was assessed using CellTiter-Glo (Promega) assay, with luminescence measured on a CLARIOStar microplate reader (BMG Labtech). For each cell line luminescence values were normalised against the value of the untreated wells.

### Telomerase activity assay

Telomerase activity was measured using the TRAPeze RT Telomerase Detection Kit (Millipore S7710). 500,000 cells were lysed using CHAPS lysis buffer, and extracts were assayed following manufacturer’s instructions, using Titanium Taq DNA Polymerase (TaKaRa). Relative telomerase activity was determined using a TSR8 standard curve in each plate, following the manufacturer’s protocol. Reactions were set up in triplicate and included heat inactivated controls within the same plate. Real-time fluorescence data was acquired using a StepOnePlus System (Thermo Fisher Scientific). The telomerase activity of 2,500-cell extract was determined from a linear plot of the log_10_ of the quantities of TSR8 control template standards versus the Ct values for their wells. The mean value the three technical replicates for each sample was calculated.

### Telo-seq

Telo-seq libraries were prepared according to the Oxford Nanopore Technologies (ONT) protocol(*42*), with in-house oligonucleotide modifications. Briefly, the Telo-Adapter and Telo-Splints were re-designed to allow multiplexing, using barcode sequences from the ONT Native Barcoding kit (Supplementary Table 1). The six Telo-Adapter oligonucleotides were annealed with the complimentary Telo-Splint, pooled for each barcode and subsequently ligated to 5 µg HMW genomic DNA. Adapter-ligated DNA was then digested with EcoRV-HF to fragment non-telomeric DNA, and treated with Klenow Fragment (3’-5’ exo-, both NEB), followed by a purification with 1x v/v AMPure XP (Beckman Coulter A63881). A second annealing step with the barcoded Telo-Splint was then followed by purification with 0.5x v/v AMPure XP, and up to six libraries were pooled for sequencing adapter ligation (SQK-NBD114.96). Pools were sequenced on either a P2 Solo or P24 sequencing device on R10.4.1 flow cells (ONT). Basecalling and demultiplexing were performed with Dorado software (ONT) using model dna_r10.4.1_e8.2_400bps_sup@v5.0.0 on Nvidia H100 hardware, using an in-house pipeline written in Nextflow. Data QC was performed with ToulligQC (https://github.com/GenomiqueENS/toulligQC). FASTQ files were aligned to the Genome assembly T2T-CHM13v2.0 [Accession ID: GCF_009914755], using ONT proprietary Telo-seq pipeline based on Schmidt. et al(*42*). Telomere length values were extracted and plots generated using R version 4.1.0 and ggplot2 version 3.5.0. Telomere length changes were calculated by plotting mean telomere length against relative PD, followed by linear regression analysis.

### Chromatin immunoprecipitation and slot blot

U2OS cells were crosslinked with 1% methanol-free formaldehyde in PBS for 10 minutes at room temperature. 0.8 M Tris pH 8 was added to quench the crosslinking for 5 minutes. Cells were then rinsed twice with PBS and subsequently harvested by scraping on ice with cold 0.05% Tween in PBS. Cells were pelleted by centrifugation and flash frozen in liquid nitrogen. For chromatin extraction, pellets of about 10 million cells were sequentially resuspended and incubated for 10 minutes in rotation at 4 °C first in Lysis Buffer 1 (50 mM HEPES pH 7.9, 140 mM NaCl, 10% glycerol, 0.5% NP-40, 0.25% Triton X-100, 1 mM EDTA) and then in Lysis Buffer 2 (10 mM HEPES pH 8, 200 mM NaCl, 1 mM EDTA, 0.5 mM EGTA). Nuclei were collected by centrifugation after both lysis steps. All buffers were supplemented with protease and phosphatase inhibitors. Pellets were resuspended in Lysis Buffer 3 (10 mM HEPES pH 8, 100 mM NaCl, 1 mM EDTA, 0.5 mM EGTA, 0.1% sodium deoxycholate, 0.5% N-lauroylsarcosine sodium salt solution) and transferred in 1 ml AFA Fiber milliTUBE (Covaris 520135). Chromatin was sheared to about 500 bp fragments with a Covaris LE220 with the following settings: duty factor 30%, peak incident power 450, cycles per burst 400, time 10 minutes. Chromatin concentration was adjusted to about 1 mg/ml. The lysate was supplemented with Triton X-100 to a final concentration of 1%, then cleared (16,000 g, 10 min at 4 °C), 50 ul of the supernatant were kept as input and 950 ul were incubated overnight at 4°C with 2 ug of SLFN11 antibody. The immunoprecipitated complexes were recovered with 50 ul of a 1:1 mixture of protein A and G Dynabeads (2 hours in rotation at 4 °C). Beads were then washed four times with wash buffer (50 mM HEPES pH 7.5, 250 mM LiCl, 0.7% Na-deoxycholate, 1% NP-40, 1 mM EDTA) and once with TE pH 8. Elution was performed with 100 ul of elution buffer (50 mM Tris-HCl pH 8, 1% SDS, 10 mM EDTA) at 65 °C for 10 minutes. DNA was recovered after incubating first with 80 ug of RNase A at 37 °C for 30 minutes, then with 80 ug of proteinase K at 55 °C overnight and purified with the QIAquick PCR purification kit. For the slot blot, the DNA was first denatured for 30 min at 55 °C with 2 volumes of 0.5 M NaOH, 1.5 M NaCl, then neutralized with 3.3 volumes of neutralizing solution (0.5 M Tris-HCl pH 7.5, 1.5 M NaCl). The DNA was then applied to a Hybond N+ positively charged nylon membrane (Amersham) and crosslinked in a 254 nm UV cross-linker with 120 mJ/cm2. The membrane was then incubated overnight with a DIG-3’ end labelled probes and processed as previously described (*70*).

### Immunofluorescence and denaturing FISH

Cells grown on 18 mm glass coverslips were fixed with 4% formaldehyde in PBS for 10 min at RT, washed with PBS three times and then processed for immunofluorescence. Cells were blocked with ADB for 1 hour at RT, incubated with primary antibodies (diluted in ADB) for 1 hour at RT, washed three times with PBS and then counterstained with Alexa Fluor conjugated secondary antibodies raised in goat (Thermo Fisher Scientific) diluted in ADB for 1 hour at RT. Cells were then washed three times with PBS and fixed again with 4% formaldehyde in PBS for 10 min at RT and then washed twice with PBS. Next, coverslips were dehydrated in 70%, 85%, and 100% (v/v) ethanol for 5 minutes each and then air-dried. Dry coverslips were hybridized with a telomeric TelC-Cy5 PNA probe (PNA Bio) in hybridizing solution (70% formamide, 0.5% blocking reagent (Roche), 10 mM Tris-HCl pH 7.5) for 90 seconds at 80°C followed by 2 hours at RT or overnight at 4°C incubation. Coverslips were washed twice with washing buffer (70% formamide, 10 mM Tris-HCl pH 7.5) for 15 min at RT. The coverslips were mounted onto glass slides with ProLong Gold antifade with DAPI (Life Technologies). Images were acquired using a Nikon Ti2 microscope fitted with a CSU-W1 spinning disk confocal unit (Yokogawa) and a Prime 95B camera (Photometrics) using Nikon CFI Apochromat LWD Lambda S 40x/1.15 WI objective lens. Following acquisition, images were imported into Fiji (NIH) for automated foci counting and colocalization quantitation.

### Terminal Restriction Fragment analysis

Terminal restriction fragment (TRF) was performed as previously described (*74*). In brief, purified genomic DNA was digested with 15U RsaI and 15U HinfI (NEB) at 37 °C, overnight. 3 μg digested gDNA was separated on a 0.5% (w/v) agarose gel overnight in 0.5x TBE buffer. Next, the gel was incubated in depurination solution (0.25 M HCl) for 30 min followed by a 30 min incubation in denaturing solution (0.5 M NaOH/1.5 M NaCl) and a 30 min incubation in neutralization solution (1.5 M NaCl/0.5 M Tris pH 7.5). The gDNA was transferred to a positively charged nylon membrane (Amersham) for 2 hours. After UV-crosslinking, the membrane was hybridised overnight at 65 °C with DIG-labelled probes as described (*74*). Chemiluminescence was detected with anti-DIG-AP antibody (Roche) and developed using CDP-star (Roche), with acquisition in a ChemiDoc MP system (Bio-Rad).

### Mouse xenograft study

All animal experimentations were undertaken in compliance with UK Home Office legislation under the Animals (Scientific Procedures) Act 1986 under project license number PP1362096 and following the ARRIVE guidelines. Animal experiments used male and female mice at 10– 14 weeks of age, bred at the Francis Crick Institute under specific pathogen-free conditions. For in vivo experiments, mice were age-matched and sex-matched where possible between groups. Randomization was undertaken to remove any biases. Each NOD.Cg-*Prkdc^scid^*/J mouse had a specific ID number and group number that did not indicate the genotype of the cells injected and treatments (vehicle/ATR inhibitor) given. All mice were handled blindly by qualified animal technicians throughout the xenograft study and subsequent analysis were also performed blindly.

5 × 10^6^ engineered eHAP iCas9 cells (WT sg*NTC*, ATRX^KO^ sg*NTC*, WT sg*SLFN11*, ATRX^KO^ sg*SLFN11*) mixed 1:1 with Matrigel (Corning) were subcutaneously implanted into the right flank of NOD.Cg-*Prkdc^scid^*/J mice. The next day all mice were given 0.2 mg/ml doxycycline hyclate in drinking water supplemented with 5% sucrose continuously for the duration of the experiment. Once tumours reached a volume between 80 and 130 mm^3^, mice were randomized into vehicle– and drug-treated groups, such that the average tumour volume was ∼100 mm^3^. AZD6738 was dissolved in DMSO to a concentration of 50 mg/ml and further diluted 1:5 in 40% propylene glycol. 25 mg/kg dose of the AZD6738 was administered every day using oral gavage for no more than 20 days. A solution of 10% DMSO and 40% propylene glycol was used as vehicle control. Tumour sizes were measured 3 times a week using a digital calliper and volumes were calculated using the formula: (length × width^2^)/2. Mice were euthanized once the mean tumour size reached <15 mm. Tumours were harvested, weighed, and measured ex-vivo at the endpoint. ¾ of each tumour were fixed in 10% formalin to perform immunohistology and ¼ snap-frozen in liquid nitrogen to assess for SLFN11 depletion.

### Immunohistochemistry of xenografts

Formalin-fixed tumours were processed and embedded in paraffin by standard techniques, sectioned at 4 μm, and then stained with hematoxylin and eosin according to standard procedures. For immunohistochemistry, samples were prepared using standard methods. In brief, tissue sections were processed for staining by microwaving in 0.01 M citrate buffer pH 6 or universal HIER antigen retrieval reagent (Abcam). After incubation with primary antibodies overnight at 4°C, samples were incubated either with biotinylated secondary antibody (Vector) followed by incubation with Avidin Biotin Complex (Vector); slides were developed in 3,3′-diaminobenzidine (DAB) substrate (Vector) and counterstained in hematoxylin, or with goat secondary antibodies followed by VectaFluor DyLight labeled Horse Anti-Goat IgG (Vector) and counterstained with DAPI. Tumours sections images were acquired at 20X using a VS200 scanner (Olympus). Stainings were quantified using QuPath software.

## Statistics

Sample number (n) indicates the number of independent biological samples in each experiment and are indicated in figure legends. Statistical details of each experiment (including the statistical tests used, exact value of n) can be found in the figure legends. Prism 9/10 (GraphPad) and RStudio were used for statistical analysis: one-way ANOVA, followed by Tukey’s multiple comparison test were used unless stated otherwise (p < 0.05 (*), p < 0.01 (**), p < 0.001 (***), p < 0.0001 (****)).

